# Uncovering a hidden diversity: optimized protocols for the extraction of bacteriophages from soil

**DOI:** 10.1101/733980

**Authors:** Pauline C. Göller, Jose M. Haro-Moreno, Francisco Rodriguez-Valera, Martin J. Loessner, Elena Gómez-Sanz

**Affiliations:** Institute of Food, Nutrition and Health, ETH Zurich, 8092 Zurich, Switzerland; Departamento de Producción Vegetal y Microbiología, Universidad Miguel Hernández, San Juan de Alicante, Spain; Área de Microbiología Molecular, Centro de Investigación Biomédica de La Rioja (CIBIR), Logroño, Spain

**Keywords:** Bacteriophage extraction, phage metaviromes, soil samples, bacterial contamination

## Abstract

**Background:** Bacteriophages are the most numerous biological entities on earth and play a crucial role in shaping microbial communities. Investigating the bacteriophage community from soil samples will shed light not only on the yet largely unknown phage diversity, but also may result in novel insights into phage biology and functioning. Unfortunately, the study of soil viromes lags far behind any other ecological model system, due to the heterogeneous soil matrix that rises major technical difficulties in the extraction process. Resolving these technical challenges and establishing a standardized extraction protocol is therefore a fundamental prerequisite for replicable results and comparative virome studies.

**Results:** We here report the optimization of protocols for extraction of bacteriophage DNA from soil preceding metagenomic analysis such that the protocol can equally be harnessed for phage isolation. As an optimization strategy, soil samples were spiked with a viral community consisting of phages from different families (10^6^ PFU/g soil): *Listeria* phage ΦA511 (Myovirus), *Staphylococcus* phage Φ2638AΔLCR (Siphovirus), and *Escherichia* phage ΦT7 (Podovirus). The efficacy of bacteriophage (i) elution, (ii) filtration, (iii) concentration, and (iv) DNA extraction methods was tested. Successful extraction routes were selected based on spiked phage recovery and low bacterial 16S rRNA gene contaminants. Natural agricultural soil viromes were then extracted with the optimized methods and shotgun sequenced. Our approach yielded sufficient amounts of inhibitor-free viral DNA for non-amplification dependent sequencing and low 16S rRNA gene contamination levels (≤ 0.2 ‰). Compared to previously published protocols, the number of bacterial read contamination was decreased by 65 %. In addition, 468 novel circularized soil phage genomes in size up to 235 kb were obtained from over 29,000 manually identified viral contigs, promising the discovery of a large, previously inaccessible viral diversity.

**Conclusion:** We have shown a dramatically enhanced extraction of the soil phage community by protocol optimization that has proven robustness in both a culture-depended as well as through metaviromic analysis. Our huge data set of manually curated soil viral contigs roughly doubles the amount of currently available soil virome data, and provide insights into the yet largely undescribed soil viral sequence space.

## Introduction

Soil bacteriophages are a vital part of soil bacterial ecology and a major reservoir of genetic material that contributes to biological evolution and diversity. They also influence the flow of nutrients as a potential cause of microbial distribution and mortality [1, 2]. Soil is known to harbour a vast abundance of phages (10^8^ −10^9^ pfu·g^−1^), with their numbers exceeding those of co-occurring bacteria by 10 – 1000 fold [3–5]. This undiscovered viral diversity could lead not only to novel findings in phage biology, but can equally promote advances in phage therapy, research for the treatment of pathogen infected humans, or agricultural cropland. Despite this ecological and medical importance, the soil virome is poorly studied compared to other ecosystems, likely due to major technical and computational limitations. Nowadays, 97 % of all viruses are thought to be found in solid matrices such as soil and sediment and adversely, only 1.8 % of all publicly available metavirome data cover those combined sources [6]. It is therefore of high importance to establish reliable methods for comparing changes in viral abundances within and across those samples. Given the physicochemical diversity of soils, the soil matrix complexity and its high microbial diversity [2], it is not surprising that no universal phage extraction protocol or standardization towards viral elution, concentration and DNA extraction have yet been proposed. Only few phage extraction protocols from soil samples have been suggested in literature, which frequently suffer from a low recovery rate of phages (< 5 %), and downstream metagenome analyses therefore require DNA amplification [1, 3, 7–17]. Moreover, the most often applied rather harsh methods may render phage isolation impossible, or imply the use of equipment that is not available in standard laboratories. Among those studies, soil phage extraction has included a wide range of elution media, such as deionized water [7, 8], SM buffer [9–11], potassium citrate buffer [4], 10 % beef extract [3, 18], modified potassium citrate buffer [1], Na/K buffer [12], or phosphate buffered saline (PBS) supplemented with beef extract [13]. Those elution media are commonly combined with mechanical approaches to disrupt phage soil interactions, such as homogenization [13], sonication [3, 4, 14], vortexing [1, 3, 11], shaking [8, 10, 15], magnetic stirring [18] or bead-beating [16, 17]. Despite these diverse approaches, the elution and recovery of soil phages remains the major bottleneck in the extraction, since more than 90 % of viruses tend to absorb to soil particles [2, 19]. Therefore, they are unintentionally removed by centrifugation or filtration techniques in the very first steps of most protocols. For phage concentration, typical techniques as tangential flow filtration [9] (TFF), or polyethylene glycol concentration [9, 10] (PEG), occasionally in combination with caesium chloride (CsCl) ultracentrifugation for purification are proposed. Those procedures, however, have only been evaluated for efficiency in other sample matrixes, such as faecal probes [20].

For functional and sequence metaviromics, purified phage DNA without contaminating bacterial or eukaryotic sequences is critical for experimental success. However, the selective extraction of viral DNA from any source, including soil matrixes, has shown to be very challenging, since most extracted phage metaviromes show bacterial DNA contamination above a proposed accepted limit (> 0.2 ‰ of ribosomal DNA reads) [8, 21]. No studies have yet assessed the influence of different phage elution, concentration and DNA extraction methods with respect to soil bacteriophages and bacterial contaminants prior to metagenomics. Here, we report the optimization of protocols for the extraction of bacteriophages from soil samples that can be used prior to metagenomics and equally be applied to infective phage particle isolation from soil. The advantage of using a culture-dependent detection method for viral recovery is evident when considering phage isolation and the obvious viability of phages used for DNA extraction. For this, soil samples were spiked with a viral community consisting of phages from different families and the efficiency of different bacteriophage elution, concentration and DNA extraction procedures were determined. Successful extraction routes were selected based on spiked phage recovery and low bacterial 16S rRNA gene contaminants. Natural agricultural soil viromes were then extracted with the optimized methods and shotgun sequenced. Our approach yielded sufficient amounts of inhibitor-free viral DNA for non-amplification based sequencing and low 16S rRNA gene contamination levels (≤ 0.2 ‰). Compared to previously published protocols, the number of bacterial read contamination could be decreased by 65 %. In total, we obtained 468 novel circularized soil phage genomes of up to 235 kb in size, from >29,000 manually curated viral contigs. This data set roughly doubles the amount of today’s available viral contigs from soil, and opens the door for discovery of a very large soil viral diversity.

## Results

### Protocol optimization strategy

The selective criteria for optimized extraction routes relied on three major parameters: (i) phage yield at each step of the extraction route, (ii) reduction of bacterial DNA contamination levels and (iii) bias minimization in the relative abundance and diversity of viruses. The efficacy of different bacteriophage elution, concentration and DNA extraction procedures was determined by monitoring spiked phage recovery by plaque assay and bacterial contamination levels by qPCR (Additional file 1: Figure S1). For this, a mock viral community consisting of phages from different families: *Listeria* phage ΦA511 (Myovirus), *Staphylococcus* phage Φ2638AΔLCR, (Siphovirus), and *Escherichia* phage ΦT7 (Podovirus), was spiked (10^6^ PFU/g soil) into agricultural soil samples. Successful extraction routes were then shotgun sequenced to gain a deeper understanding of the soil viral community and to compare the soil viral diversity in each extraction route, including data generated from a literature-based approach.

### Resuspension of soil phages

A proper suspension of bacteriophages from soil particles is crucial to retain viral diversity and reproducibility. We compared the most promising elution buffers found in literature to maximize bacteriophage suspension from agricultural soil samples (Figure 1). For this, the most commonly used (or elsewhere optimized) elution buffers such as SM buffer [9, 10], amended potassium citrate (AKC) [1], 10 % beef extract [3, 18] or PBS amended with beef extract [13] were assessed and compared in viral yield using plaque assay. When soil samples were suspended using SM or AKC buffer, as little as 0.5 to 5 % of all spiked phages could be recovered (Figure 2a and 2b). Those simple salt-supplemented buffers, however, provided good filtration properties after soil suspension and did not interfere with any downstream analysis. Protein-supplemented elution buffers, such as PBS + 2.5 % beef extract or 10 % beef extract, recovered a compelling number of spiked bacteriophages (51.1 % and 66.7 %, respectively). Yet, when performing phage elution protocols with more than 300 g of soil, any buffer that contained beef extract resulted to be a poor choice. Vacuum-filtration attempts with filter pore size smaller than 1 µm were instantly clogged and techniques applied downstream (qPCR, microscopy or concentration methods) failed completely. It is therefore evident that beef extract is a very efficient supplement to suspend soil phages, but it equally dissolves other organic compounds that interfere with ensuing techniques.

**Figure 1.**
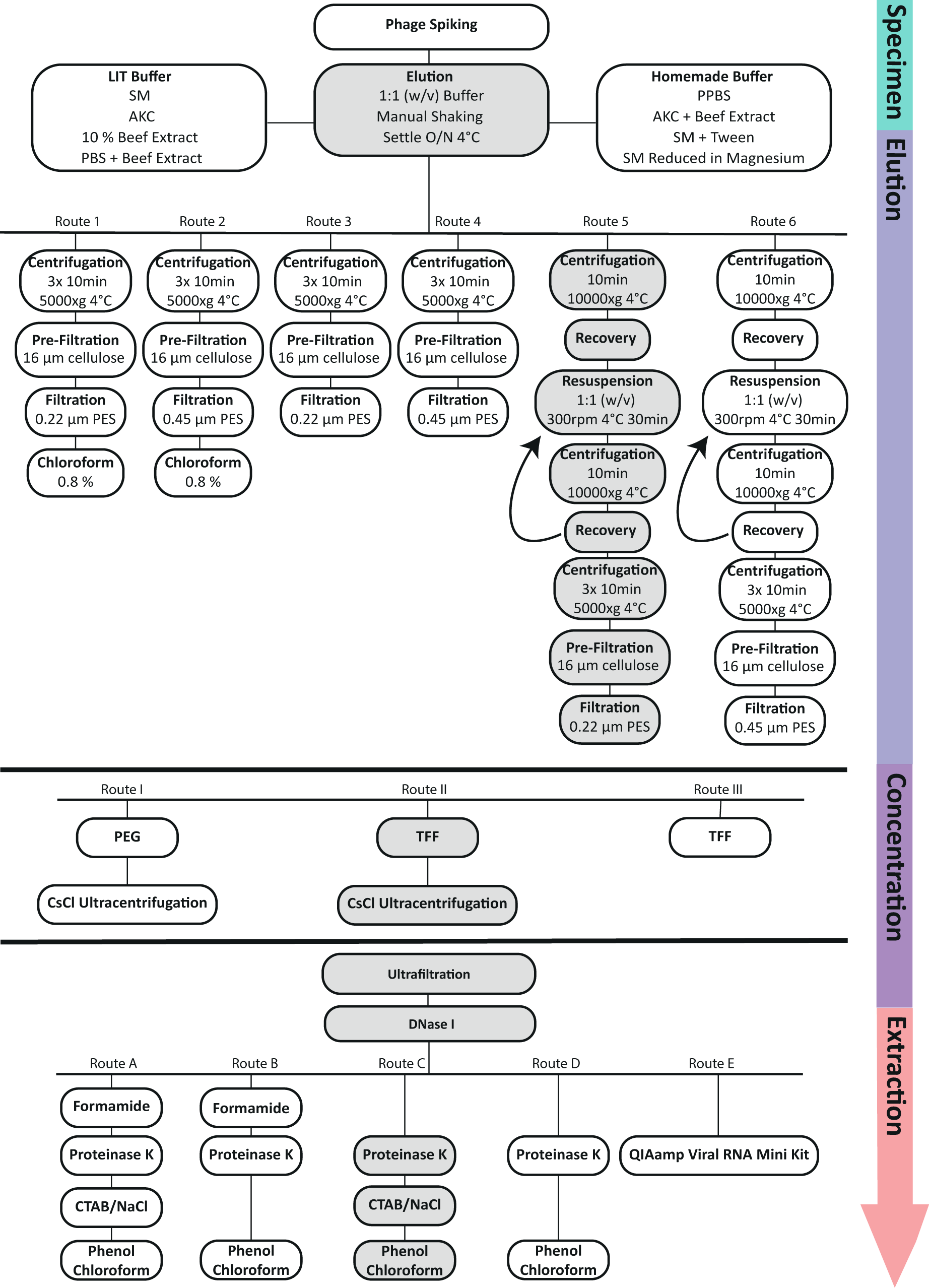
Optimization protocols for bacteriophage elution and extraction. Optimization strategy for phage DNA extraction from soil samples prior metagenomic analysis. An efficient concentration and purification of soil bacteriophages together with a complete removal of bacterial DNA is aimed. Bacteriophage elution (Route 1-6), concentration (Route I-III), and DNA extraction (Route A-E) methods are evaluated for efficiency using a spiked soil sample (ΦA511, Φ2638AΔLCR, ΦT7 at 10^6^ PFU/g soil) and plaque assays. Bacterial DNA depletion was assessed using 16S rRNA qPCR. Optimal phage extraction route is highlighted in grey as verified by sequencing metagenomics.

**Figure 2.**
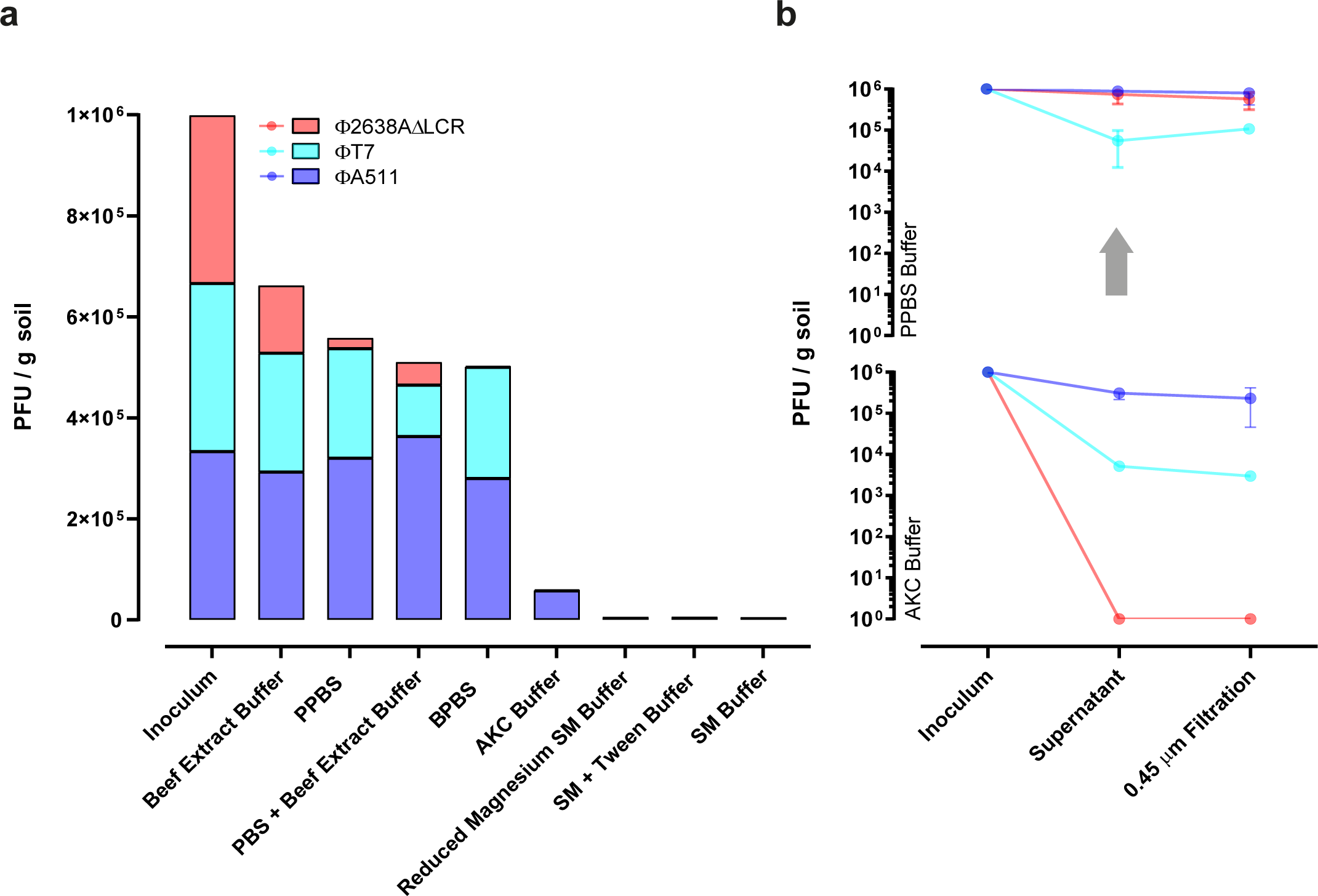
Bacteriophage elution buffers for soil samples. Bacteriophage elution from soil samples was assessed with different elution buffers. The ability to elute phages from soil samples was tested for elution buffers previously published in literature or novel constituted elution buffer. Soil samples were spiked with an artificial viral community (ΦA511, Φ2638AΔLCR, ΦT7 at 10^6^ PFU/g soil) and bacteriophage yield was assessed using plaque assay. (**a**) SM buffer [9, 10]; 10 % beef extract buffer [3, 18]; PBS supplemented with beef extract [32]; AKC [1]. (**b**) Elution amelioration measured by plaque assay (Mean ± SD) when using optimized PPBS instead of literature proposed elution buffer.

Our elution buffers were, therefore, adapted and assessed for efficiency with the following modifications: SM buffer was supplemented with 0.01 % tween or reduced in magnesium, and additionally, two novel in-house elution buffers were designed consisting of PBS with either a BSA (PPBS) or beef extract (BPBS) supplementation. For PPBS (protein supplemented PBS), the 10 % PBS, 1% KC and 150 mM MgSO_4_ were adapted from the optimized AKC buffer [1], while EDTA was removed and substituted with a 2 % BSA supplementation (Figure 1). For BPBS (beef extract supplemented PBS), BSA from PPBS was replaced by beef extract. Salt components in these elution buffers serve as pH and viral particle stabiliser [1], whereas the protein addition as BSA or beef extract offers viral binding sites and disrupts soil-viral interactions [18]. The adjusted SM buffer failed to improve recovery and, hence, did not prevent non-specific phage soil interactions. PPBS performed only negligibly worse in the elution of bacteriophages compared to 10 % beef extract (55.8 % versus 66.2 % recovery, respectively), and did not trigger any of the technical difficulties described above, even when applied to large samples (>1 kg soil). Interestingly, when replacing BSA with beef extract (BPBS) in this optimized buffer, no enhanced phage recovery was observed. As previously suggested [4, 12], recovery of spiked bacteriophages was further optimized by resuspending the soil pellet thrice (Figure 1, Route 5-6), resulting in a relative increase in recovery of 48 % compared to a single suspension step (45.7 % and 67.3 % recovery, respectively). Our optimized elution protocol allows, hence, a virtually complete recovery of infective bacteriophages with simple methods such as adjusting elution buffer constitution and washing rounds of the soil pellet (Figure 2b). In summary, soil samples should be eluted in equal volumes of PPBS, manually shaken by inversion and let to settle overnight at 4 °C. The suspended soil should then be centrifuged as described in previous sections, the supernatant kept aside and the pellet resuspended in another volume of PPBS for a total of three rounds [4, 12].

### Removal of bacterial contaminants using centrifugation and filtration

After elution optimization of bacteriophages from soil particles, the complete removal of contaminating bacterial cells and thus bacterial DNA was attempted. Several techniques previously described in literature, such as centrifugation steps prior to 0.8 µm [22], 0.45 µm [1, 17, 23] or 0.22 µm [3, 9, 10, 12, 15, 24–26] PES filtration (Figure 1, Route 3-6), some coupled with a chloroform treatment [9, 12] (Figure 1, Route 1-2) were assessed. The benefit of using PES as the filter material was already established elsewhere and applied here [22]. Any filtration attempt with pore size < 16 µm was impaired if the single (Figure 1, Route 1-4) or mixed (Figure 1, Route 5-6) supernatants were not centrifuged thrice at 5,000 x g for 10 minutes. Filtration procedures and the spiked phage community were measurably not impaired when using low-speed centrifugation to remove impurities. As summarized above, published protocols in the literature suggest either a 0.22 µm, 0.45 µm or 0.8 µm filtration of extracted metaviromes to decrease bacterial contamination below a reasonable threshold and simultaneously not impair viral yield or diversity. A reductive effect on soil phage diversity when using a 0.22 µm filter to decrease external contamination, has however not been evaluated yet. In this optimization protocol, no significant difference in spiked phage recovery was observed when comparing a 0.22 µm to 0.45 µm filter pore size (unpaired t-test, p-value=0.7782, n=9) (Figure 3a). In addition, a 16S rRNA qPCR analysis revealed that both 0.45 µm and 0.22 µm PES filtration techniques removed more than 99.9 % of all bacterial 16S rRNA genes. The 0.22 µm filtration, however, decreased bacterial gene contamination significantly better (unpaired t-test, p-value <0.0001, n=6) (Figure 3b). A higher recovery of phages in metaviromic samples was recently reported when substituting a 0.22 µm with a 0.45 µm filtration step [23]. Along these lines, the 0.45 µm filtration was nevertheless chosen here for further optimization purposes in order to avoid a potential bias in the native soil viral community. Both routes (0.22 and 0.45 µm filtration) were then selected for shotgun sequencing analysis of soil metaviromes to provide definite answers to bacterial contamination levels and viral diversity in each approach (see below).

**Figure 3.**
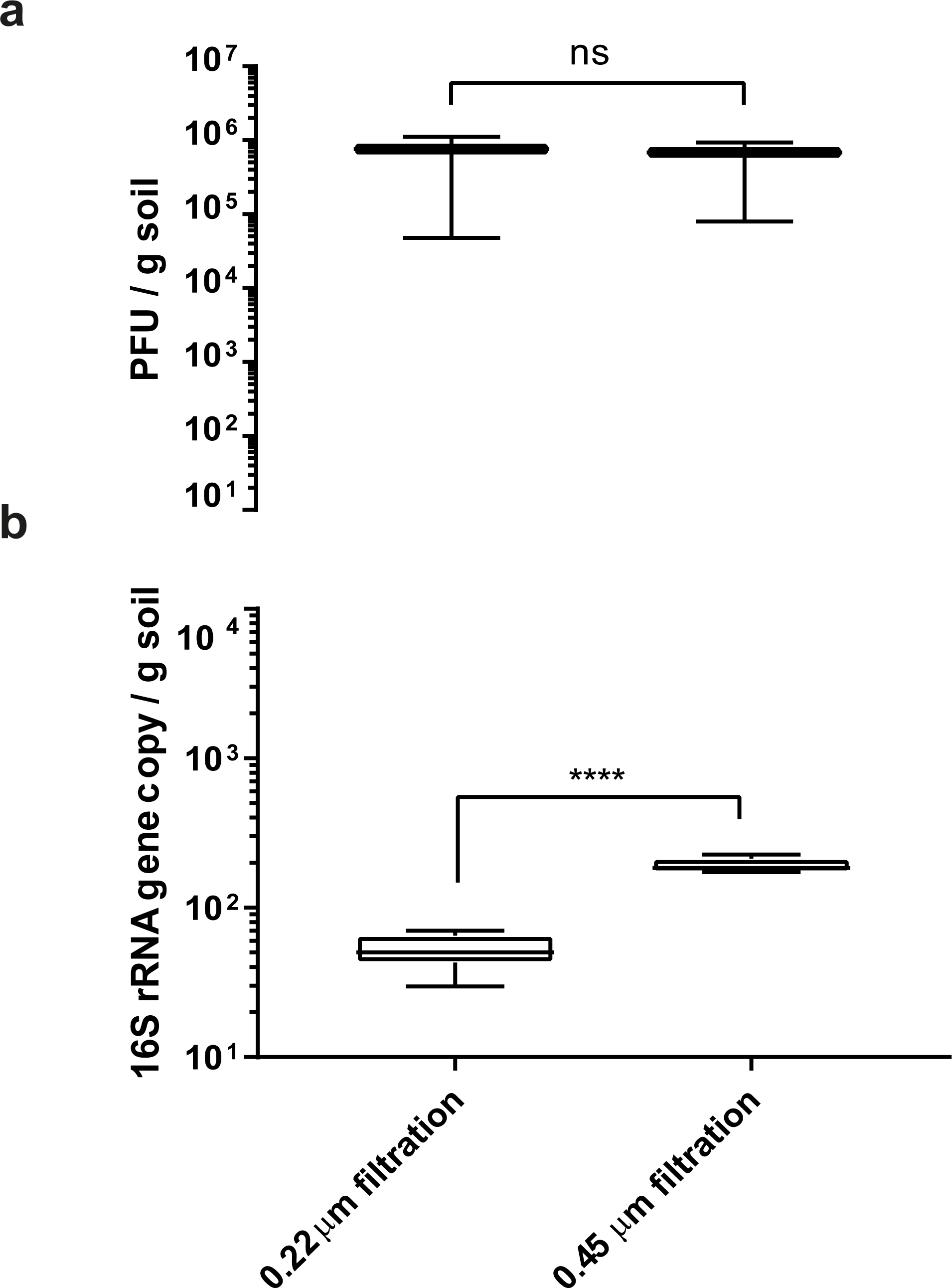
Effect of filtration pore sizes on bacteriophage recovery and bacterial contamination. Recovery of spiked bacteriophages (a) and respective bacterial DNA contamination (b) with a 0.22 µm or 0.45 µm PES filtration. Spiked bacteriophages were detected after bacteriophage elution and filtration of the soil suspension using plaque assay. Bacterial DNA contamination was assessed using Taqman qPCR amplifying the 16S rRNA gene. Boxplot and whiskers (min to max) with ns equals p=0.7782 (n=9), **** equals p<0.0001 (n=6).

Besides centrifugation and filtration, the efficiency of chloroform treatment to remove bacterial contamination was assessed (Figure 1, Route 1-2). Chloroform treatment is generally a rather impractical approach, because both bacteriophages in the environmental sample [9], as well as downstream concentration devices, are sensitive to chloroform. A maximum concentration of 0.8 % chloroform is supported when using a PES tangential – or regular filtration, which in turn did not reduce bacterial DNA contamination (data not shown). A chloroform treatment prior to phage concentration was therefore excluded from the protocol.

### Concentration of viral particles from soil samples prior to DNA extraction

PEG and TFF concentration techniques in combination with ultrafiltration are commonly used techniques to concentrate viral particles from large volumes. These techniques have been described in detail elsewhere [9, 10, 20, 27] and were assessed here in eluted soil samples. An optimal concentration technique should concentrate viral particles without introducing a bias to the native viral community, and equally reduce the suspensions’ volume sufficiently to allow DNA extraction. Spiked phages were eluted from soil samples using the optimized elution protocol, filtrated through a 0.45 µm PES filter and subjected to PEG or TFF (Figure 1, Route I-II). As shown in Figure 4, both concentration techniques performed equally good in concentrating spiked phages and no differences in spiked phage yield after concentration was observed. Furthermore, viral suspensions were reduced in both approaches to a final volume of 20-50 ml, which allowed CsCl ultracentrifugation and ultrafiltration. Like in the case of filtration, the spike dependent approach for evaluating efficiency of those concentration methods, has not resulted in a clear conclusion. Both routes were therefore selected for shotgun sequencing to elucidate which route revealed superior by obtaining highest viral recovery and diversity (see below). CsCl ultracentrifugation concentrates and purifies phages from samples or pure cultures. It is however not only technically demanding, but also cost and equipment intensive. The necessity of a CsCl purification for soil samples prior to DNA extraction was evaluated by extracting soil viruses with and without CsCl centrifugation prior ultrafiltration (Figure 1, Route II and III). Ultrafiltration units were coated with PBS + 1 % BSA to reduce viral absorption as suggested and optimized elsewhere [27]. Purification of soil samples with CsCl ultracentrifugation seemed to slightly diminish spiked viral yields (Figure 4a and 4b). However, this loss in PFU could also be attributed to a loss in infectivity rather than a loss in viral yield caused by centrifugation conditions [20]. When avoiding a CsCl purification step prior to ultrafiltration (Figure 1, Route III, and Figure 4c), the centrifugal filters clogged while concentrating and the remaining volume could not be reduced to less than 5 ml. A purification of soil viral suspensions using CsCl ultracentrifugation was therefore implemented in the optimized protocol, which ensured a concentration to a final volume below 300 µL. When aiming for isolating infective viral particles from soil samples, however, this purification can easily be omitted.

**Figure 4.**
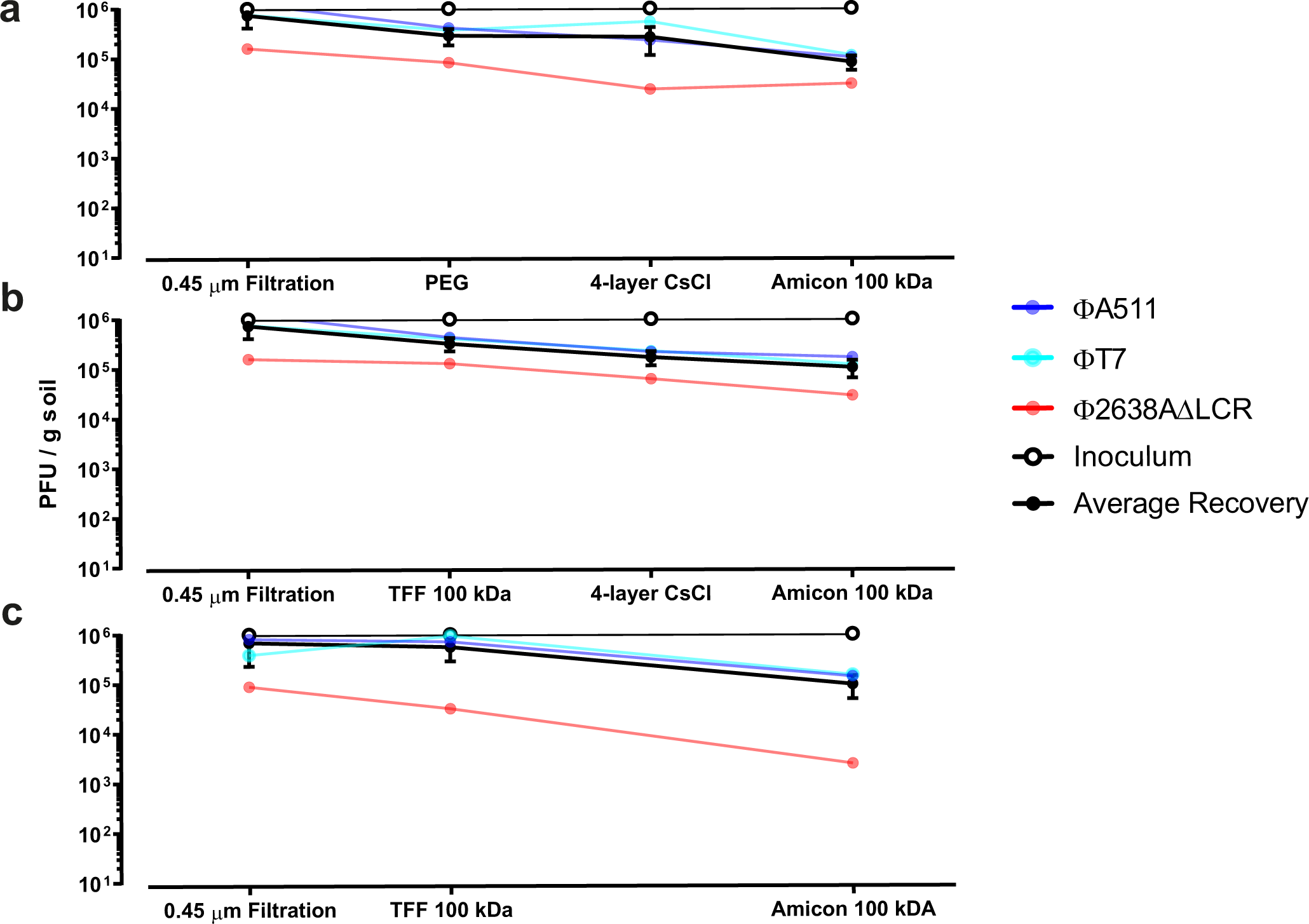
Recovery of spiked bacteriophages. Spiked bacteriophage recovery monitored in concentration (PEG, TFF, and ultrafiltration) and purification methods (CsCl ultracentrifugation). Soil samples were spiked with an artificial viral community (ΦA511, Φ2638AΔLCR, ΦT7 at 10^6^ PFU/g soil), phages were eluted with the optimized protocol and subjected to concentration protocols. A reduction in PFU at a given technique quantifies (Mean ± SEM in triplicates) the loss of spiked bacteriophages.

### Phage DNA extraction from soil samples

Ultrafiltrated concentrated samples were treated with DNase I to remove free DNA that is not enclosed by viral capsids, and brought up to a final volume of 600 µl. From here, phage DNA was either extracted using modified phenol/chloroform extraction routes (Figure 1, Route A-D) or QIAamp Viral RNA Mini Kit (Qiagen) according to manufacture instructions (Figure 1, Route E). DNA extraction using the suggested kit resumed in a > 10 fold reduction of viral DNA than other proposed DNA extraction routes, and was therefore excluded from the optimized protocol (data not shown). The phenol/chloroform DNA extraction was adapted from Thurber et al. [9], whereas the necessity of formamide and CTAB in both viral DNA extraction routes was assessed by bypassing these steps separately or in combination. DNA extraction routes with larger volumes that originated directly from the TFF concentrated sample (Figure 1, Route III, no CsCl purification), resulted in a jellification of the sample after the addition of formamide, or presented a contamination with qPCR inhibitors that even flawed DNA measurements if formamide was neglected. Extraction routes covering a formamide treatment resulted therefore either in a complete loss of metaviromic DNA through a jellification of the sample (Figure 1, Route III), or reduction in DNA yield from CsCl purified routes (Additional file 3: Table S2). Formamide treatment is thus decreasing the viral DNA yield irrespectively of purification and was therefore excluded. A CTAB/NaCl treatment on the other hand, did not correlate consistently with a disadvantageous outcome and was further highly dependent on the experimental setup. For the sake of simplicity, a CTAB/NaCl treatment of viral suspensions was hence incorporated in viral DNA extraction (Additional file 3: Table S2). When performing the optimized protocol, a sufficient amount of pure viral DNA was extracted from 400 – 1000 g of soil (Additional file 3: Table S2, Table 1), which allowed direct sequencing without using multiple displacement amplification.

**Table 1.** DNA yield, trimmed reads and assembled contigs from optimized phage extraction routes.

| Sample | DNA (ng/μl) | Trimmed Reads (Mill) | % of Reads Surviving | Contigs (> 5 kb) | Average kb | Nucleotides Assembled (Mb) |
| --- | --- | --- | --- | --- | --- | --- |
| 0.22 μm TFF | 2.7 | 104 | 94.2 | 17,698 | 11.22 | 196.5 |
| 0.22 μm PEG | 1.70 | 60 | 92.4 | 7,583 | 11.15 | 84.5 |
| 0.45 μm TFF | 56.6 | 83 | 91.0 | 12,453 | 11.58 | 164.0 |
| 0.45 μm PEG | 2.44 | 64 | 92.9 | 10,493 | 13.17 | 121.4 |

### Metagenomic analysis

By using a spiked phage community as reporter, it was not possible to provide definite answers regarding an optimal filter pore size (0.22 μm vs 0.45 μm) or soil phage concentration method (PEG vs TFF). The four optimized phage DNA extraction routes (0.22 μm + TFF, 0.22 μm + PEG, 0.45 μm + TFF and 0.45 μm + PEG) were hence compared for viral richness, diversity and bacterial DNA contamination levels using metagenomic analysis. Viral DNA was extracted from 1 Kg of soil for each extraction route and paired-end shotgun Illumina sequenced with 76 million reads per route. Over 90 % of all raw reads survived the initial data pre-processing as trimming and size exclusion and a total of 311 million reads from all four metaviromes remained. Those reads were assembled into 48,227 contigs (> 5 kb) with average lengths between 11.11 and 13.17 kb (Table 1). As a normalization measure and to exclude a potential direct effect of the number of reads to the assembled contigs, a sub-assembly with 60 million reads for each sequenced extraction route was performed (Additional file 4: Table S3). This sub-assembly resulted in less assembled contigs for each metavirome, indicating an incomplete coverage of the metaviromic diversity. The abundance of contigs per metavirome, however, deceased proportionally such that the 0.22 μm + TFF metavirome still displayed the highest amount of assembled contigs independently from the number of reads used for assembly.

To appraise bacterial contamination levels, the percentage of 16S rRNA reads in each method was assessed based on confirmed 16S rRNA reads after ssu-align, and were taxonomically classified by sequence match against the RDP [28]. Contamination of 16S rRNA genes in both 0.22 μm filtrated metaviromes was below 0.2 ‰ (0.019 % in PEG and 0.018 % in TFF, respectively). Bacterial DNA contamination in 0.45 μm filtrated metaviromes was significantly higher and above the suggested threshold for metaviromes of 0.2 ‰ (0.054 % in PEG and 0.065 % in TFF, respectively) [21] (Figure 5). Similarly, the LIT established metavirome solely filtrated through a 0.22 μm PES filter resulted in equally high contamination levels (0.053 %) (Figure 5). Our proposed optimized protocol resulted, therefore, in a high reduction of external bacterial DNA contamination (> 65 %) compared to previously published protocols. Interestingly, each metavirome, regardless of the phage DNA extraction route, had *Candidatus Saccharibacteria* as the predominant contaminating bacterial phylum (Figure 5). *Saccharibacteria* are exceptionally small Gram-positive cocci (200 - 300 nm) that are able to pass through even a 0.22 μm filter due to their size [29]. Those bacteria are thus being concentrated along with bacteriophages in all protocols, and their DNA extracted alongside. In the 0.22 μm filtrated metaviromes, *Saccharibacteria* composed > 60 % of all contaminants, which decreased to 40 % in the 0.45 μm filtrated viromes. However, this decrease should not be mistaken for an absolute value, as 0.45 μm filtrated filtrated viromes displayed two to three times higher external contamination (Figure 5).

**Figure 5.**
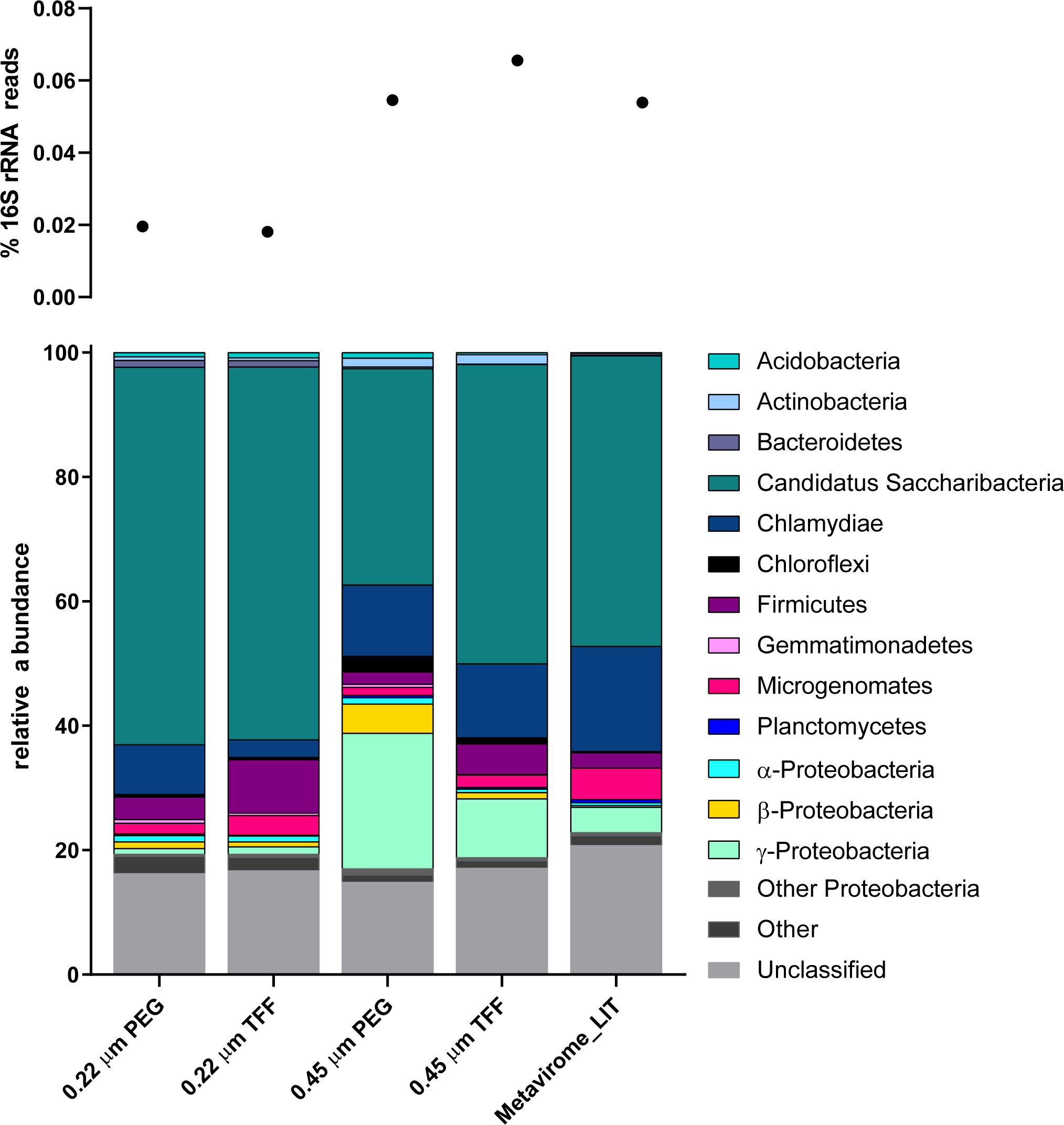
16S rRNA gene contamination. 16S rRNA gene contamination in the five extracted and sequenced metaviromes and one metagenome. Metaviromes filtrated with a 0.22 μm filter and extracted with the optimized protocol are below the previously established threshold for sufficiently purified metaviromes [21] (a). *Candidatus Saccharibacteria* are the predominant contaminant of all metaviromes irrespective of the extraction method applied (b).

After assembly, the 48,227 remaining contigs were manually inspected and classified as virus, if viral hallmark genes such as terminases or structural proteins were present. Equally, those contigs that harboured bacterial genes such as ribosomal sequences were separated from the viral fraction and classified as bacteria. After identification by manual curation, 13,114 contigs were classified as virus and another 13,519 as bacteria, whereas 21,586 remained unclassified due to insufficient annotation (hypothetical proteins or none) (Table 2). Manually classified viral contigs from each metavirome were then pooled and redundant contigs (clustered at > 99 % identity) were removed. This initial clustering analysis resulted in 10,886 (74 %) unique and partially complete viral genomes from all four extracted soil metaviromes. On the basis of overlapping ends (more than 10 bp), we could extract 379 novel, circularized phage genomes with lengths from 5.1– 235 kb (average 58.9 kb) from this non-redundant viral fraction. In addition, 89 complete potential phage genomes (sizes between 5 – 70.9 kb) were identified in contigs left unclassified (Table 3).

**Table 2.** Manually assigned viral, bacterial and unclassified contigs for each sequenced metavirome.

| Sample | Viral Contigs | Bacterial Contigs | Unclassified Contigs |
| --- | --- | --- | --- |
| 0.22 μm TFF | 6,345 | 1,549 | 9,801 |
| 0.22 μm PEG | 2,453 | 1,070 | 4,058 |
| 0.45 μm TFF | 2,092 | 7,067 | 3,286 |
| 0.45 μm PEG | 2,224 | 3,833 | 4,441 |

**Table 3.** Metaviromic contigs after removal of redundant contigs with clustering analysis I and II.

|  | Clustering I | Circularized | Size of Circularized Phage Genomes | Clustering II |
| --- | --- | --- | --- | --- |
| **Viral Contigs** | 10,886 | 379 | 235 – 5.1 kb | 8,835 |
| **Bacterial Contigs** | 10,645 | - | - | 8,349 |
| **Unclassified Contigs** | 14,180 | 89 | 70.9 – 5 kb | 12,520 |
| **Total** | 35,710 (74.0 %) | 468 | 235 – 5 kb | 29,704 (61.7 %) |

Viral diversity in each extraction route was compared by removing all redundant viral contigs with clustering 1 and 2 information leaving 29,704 (61.7 %) contigs classified as either viral, bacterial or unknown origin (Table 3). A subset of 20 million reads from each sequenced metavirome was then separately mapped against this manually curated and trimmed viral community to estimate viral recruitment in each extraction method. In the 0.22 μm filtrated metaviromes, 97.5 % of all recovered viruses were present in the TFF route (967 unique viral contigs > 5 kb), whereas 88.9 % were recovered by PEG concentration (219 unique viral contigs > 5 kb) (Figure 6a). Similarly, by comparing TFF versus PEG concentration in the 0.45 μm filtrated metaviromes, it is evident that TFF performed better by recovering a higher percentage of unique soil viruses (Figure 6b). Particular soil viruses exclusively detected in the TFF route infected predominantly the phylum *Actinobacteria* (43 %), whereas 41 % could not be identified due to lack of annotation. In contrast, only 2 % of the unique viruses recovered by PEG infected *Actinobacteria* and as many as 77 % could not be characterized due the low number of functional gene predictions or lack of signature genes.

**Figure 6.**
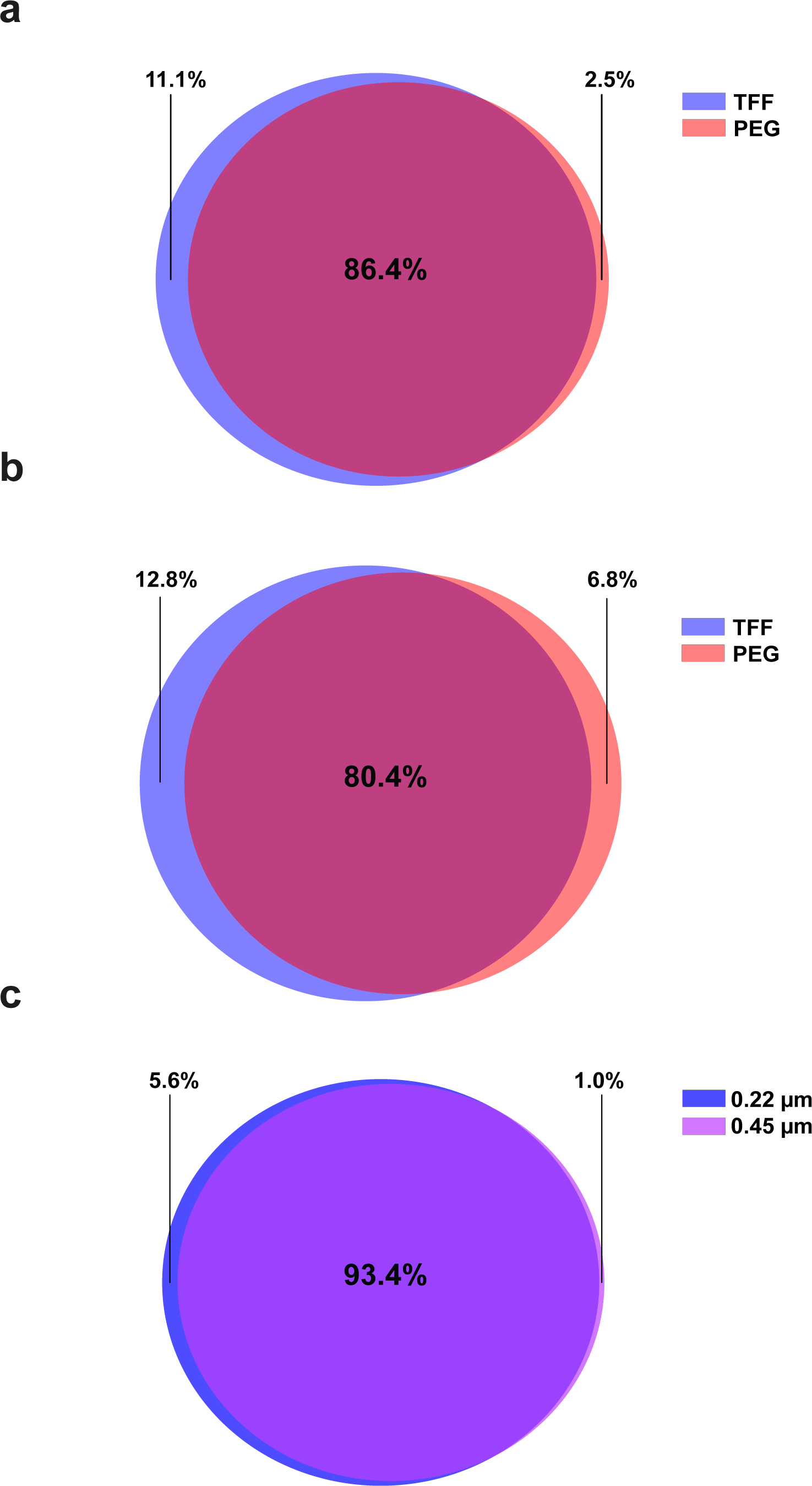
Native soil viral diversity. Native soil viral diversity in each optimized phage DNA extraction route. (a) Viral diversity in the 0.22 μm filtrated metaviromes. (b) Viral diversity in the 0.45 μm filtrated metaviromes. (c) Viral diversity in TFF concentrated metaviromes, filtrated through either 0.22 μm or 0.45 μm PES filter.

Metaviromes filtrated with a 0.45 μm filter (0.06 % 16S rRNA reads) instead of 0.22 μm (0.018 % 16S rRNA reads) harbour 70 % more bacterial DNA contamination (Figure 5), and also failed to recover unique soil viruses. Indeed, the 0.45 μm filtrated metaviromes recovered 94.4 % of all viral contigs and therefore consist of less unique soil viruses compared to a 0.22 μm filtrated virome (99 % recovery) (Figure 6c). In addition to viral diversity, variations in the percentage of reads that matched to viral, bacterial or unknown contigs were observed. The percentage of reads that matched with bacterial contigs extracted from 0.22 μm and 0.45 μm filtrated optimized protocols were 4.2 % and 15.1 %, respectively, confirming the reduced bacterial contamination in the 0.22 μm filtrated metaviromes. In addition, those metaviromes displayed the highest sequence affiliation to viral contigs, recruiting more than 25 % of the reads. Recruitment of viral reads decreased to less than 15 % in 0.45 μm extracted metaviromes (Figure 7). Notably, the recruitment rates for the unclassified fraction was considerably higher in 0.22 μm filtrated samples.

**Figure 7.**
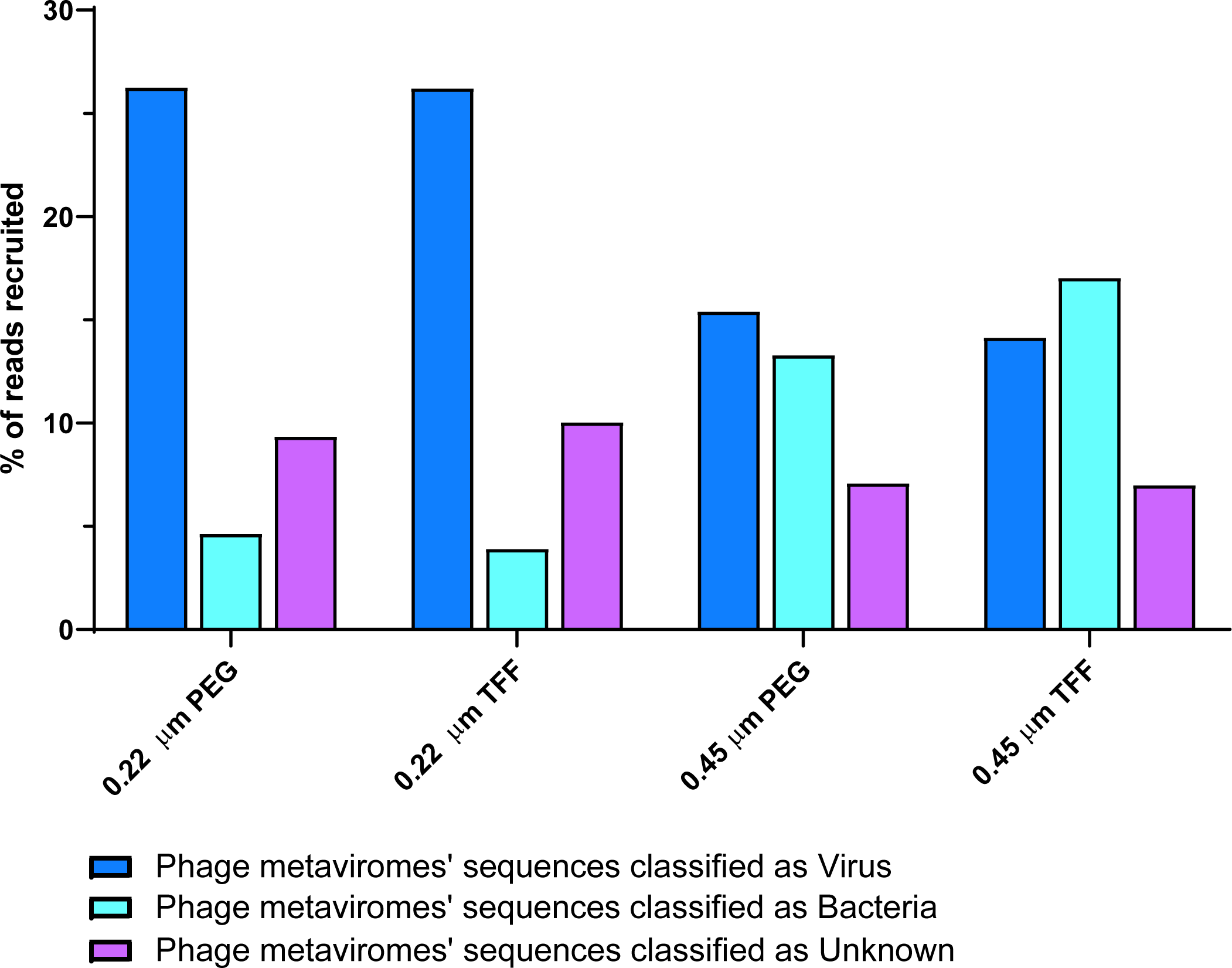
Metavirome read association. Metavirome reads associated to manually classified viral, bacterial or unknown contigs from the five sequenced metaviromes and one metagenome.

## Discussion

The study of soil viromes lag far behind any other ecological model system, due to the heterogeneous soil matrix that rise major technical difficulties in the extraction process [2]. Resolving these technical challenges and establishing a standardized extraction protocol is therefore a fundamental prerequisite for replicable results and comparative virome studies. We here report the optimization of protocols for extraction of bacteriophage DNA from soil preceding metagenomic analysis, such that the protocol can also be harnessed equally well for phage isolation. As anticipated, the elution of virus particles from soil samples has crystallized to be the major bottleneck in the present study [1, 2], due to the fact that > 90 % of bacteriophages tend to absorb to soil particles [19]. Indeed, all suggested elution buffers in the literature [1, 3, 9, 13, 18, 20] were either found to perform insufficiently in the recovery of bacteriophages from soil (< 5 % of spiked phages), or resulted in major technical limitations due to complete inhibition of downstream filtration procedures. We designed an optimal elution buffer PPBS, consisting of ionic salt compounds supplemented with 2 % BSA that disrupts phage soil particle interactions through competing for viral binding sites. This finding is consistent with Lasobras et al. [18], who reported that an optimal elution of virus particles from soil requires either a proteinaceous material that competes for viral binding sites or chaotropic agents which alter the favourability of absorption. The beneficial action observed with PPBS buffer was not enhanced when substituting BSA with beef extract, which supports the favourable effect of BSA in viral elution and validates BSA as equally, if not superior, to a beef extract supplement. In summary, the optimized elution protocol described here results in a recovery of virtually all spiked bacteriophages whereas no technical difficult or harsh method was applied to maximise viral recovery. As a major advantage, this gentle optimized elution protocol allows the isolation of infective viral particles through omitting techniques that could result in tail breakages or defective particles.

For shotgun sequencing or functional metagenomics, a maximum reduction of bacterial DNA contamination is crucial to allow data analysis. Filtration of unknown viral suspensions to remove bacterial contamination is, therefore, an extensively discussed topic in previous studies. Published protocols in literature suggest either a 0.22 µm, 0.45 µm or 0.8 µm filtration of extracted metaviromes to decrease bacterial contamination below a threshold and simultaneously not impair viral yield or diversity. Recently, some optimization studies highlighted the beneficial action of 0.45 µm filter and reported a significantly higher viral yield when substituting 0.22 µm with 0.45 µm filtration [23]. A compromising effect of soil phage communities when using 0.22 µm filter pore size had however not been evaluated yet. As shown by shotgun sequencing of 0.22 µm and 0.45 µm filtrated metaviromes, a 0.22 µm PES filter removed more bacterial DNA contamination compared to a 0.45 µm PES filter, while simultaneously did not compromise total viral recovery and diversity from soil samples. Indeed, less viral diversity was observed in metaviromes filtrated through a 0.45 µm pore-size, which is probably due to the increased bacterial DNA contamination and thus, an impaired assembly of viral contigs (Figure 6). This finding is supported by our recruitment analysis, which revealed similar amounts of reads recruited to the manually classified viral or bacterial fraction (14.7 % and 15% respectively) in 0.45 µm metaviromes. The vast majority of reads recruited in the 0.22 µm metaviromes however, matched to viral contigs and < 5 % to bacterial contigs (Figure 7). The 0.45 µm metaviromes suffered thus from more bacterial contamination, which does not only complicate approaches such as functional metagenomics, but also hinders sequencing analysis due to an impaired assembly. Using the proposed optimized protocol for elution and filtration of soil viruses, the 16S rRNA gene contamination could be reduced to a level below the recommended threshold of 0.2 ‰ [21]. This finding is consistent with Castro et al. [20], who observed a considerably lower host DNA contamination when relying on both centrifugation (e.g thrice 5000 x g) and filtration techniques. Interestingly, the predominant bacterial contaminant in all sequenced metaviromes was found to be *Candidatus Saccharibacteria*. Due to their size, *Saccharibacteria* can pass untroubled through even a 0.22 μm filter and are thus being concentrated along with the bacteriophages in any protocol. Unfortunately, those bacteria are not only found in soil, but in many other environmental samples such as sludge and activated sludge from wastewater treatment plants, human saliva and the gut microbiome [29]. Any metavirome extracted from those samples must, therefore, be either manually curated to remove bacterial reads, or handled with great care to reach valid conclusions.

In order to concentrate bacteriophages from large suspension volumes, most commonly used approaches as TFF and PEG precipitation were compared here. Independently from the filtration technique applied, TFF performed better in recovering and concentrating soil phages and, therefore, revealed a greater soil viral diversity. After concentration, a purification of soil viral suspensions using CsCl ultracentrifugation was implemented in the optimized protocol before viral DNA extraction to remove inhibitors and allow ultrafiltration concentration to a finale volume below 300 µL. As resolved here, CsCl ultracentrifugation needs to be applied for purification purposes prior to DNA extraction, but can easily be omitted when aiming for the isolation of infective viral particles. In this study, viral DNA extraction was carried out as described by Thurber et al. [9], whereas a shortening of the protocol by omitting a formamide treatment was implemented. The formamide treatment did not improve DNA extraction regardless of CsCl purification, and resulted in either a jellification (no CsCl purified sample) or a reduction in the total DNA yield (CsCl purified sample). The optimized DNA extraction protocol therefore omits unnecessary or detrimental steps found in the literature and shortens the protocol by one day.

After Illumina sequencing of our four extracted metaviromes, 48,227 contigs were assembled, manually screened and classified as either viral, bacterial or unknown sequences according to signature genes and annotation. With this manual classification, we confirmed 10,886 contigs as unique partial viral genomes and another 14,180 as putative viral contigs without bacterial hallmark genes. This finding more than doubles the currently 10,009 published viral contigs from soil viromes [30], and highlights the still very fragmented nature of available datasets from soil ecosystems. Out of our manually identified viral fraction, 379 novel and complete phage genomes in size up to 235 kb were extracted, which average size is consistent with the presently isolated dsDNA viruses [31]. In addition, we obtained another 89 closed and novel putative phage genomes in length up to 70.9 kb from the contigs classified as unknown. This shear number is even more noticeable when compared to the recently 999 extracted complete viral genomes (> 100 bp) from over 125,000 contigs that derived of 3,042 geographically diverse environmental samples [31]. In summary, our newly developed protocol for the extraction of soil bacteriophages has proven to produce robust results not only in a culture dependent analysis through spiked bacteriophages, but also through sequencing metaviromics and promises exciting insights into the immense viral diversity of the previously largely inaccessible soil virome.

## Conclusion

To our knowledge, this is the first optimized bacteriophage extraction protocol for soil, which allows sequencing analysis and infective phage isolation. We have shown a dramatically enhanced extraction of the soil phage community by protocol optimization and present soil metaviromes harbouring 468 novel and uncharacterized putative soil bacteriophages. Our huge data set of manually curated soil viral contigs provides insights into the yet largely undescribed soil viral sequence space and will double the amount of currently available soil virome data. Our optimized protocol has proven robustness in both a culture-depended and metaviromic analysis, and could be valuable for any viral extraction from solid matrixes. The only measure that needs to be undertaken before application is the development of an adequate elution buffer that suspends viral particle from the respective sample. Our guidelines for elution buffer design will facilitate such proposals in the future. These findings, in coordination with technical advances in sequencing metagenomics and phage biology, will enable a better understanding of viral diversity and phage ecology in the future.

## Methods

### Soil samples

Agricultural soil samples were collected from a long-term soil experimental field (ZOFE, Zurich Organic Fertilization Experiment), located in a rural area surrounding Zurich (Agroscope, Reckenholz, Switzerland). Soil consisted of 56 % sand, 28 % silt and 14 % clay (in mass %: 0.6 soil organic carbon, 1.1 soil humus, pH ~5.7). Soil samples for spiking and optimization of extraction routes were taken in September 2017(replica I-V, unfertilized control samples), while those for sequencing metagenomics were taken in October 2018 (replica I-V, farmyard manure fertilized samples). For each replica plot, the first 10 cm of superficial soil was sampled randomly six times with equal distances apart, and the upper rhizosphere (2-3 cm) removed. Samples were passed through a 2 mm pore size sieve. Replica plots were combined in batches of 400 g or 1,000 g, and stored immediately at -80 °C. Prior usage, the soil was defrosted at 4 °C for 4 hours.

### Design of viral mock-community

For accurate phage quantification, three strictly lytic dsDNA phages were propagated and spiked at a concentration of 1 × 10^6^ pfu/g soil: ΦA511 (*Myoviridae*), Φ2638AΔLCR (*Siphoviridae*) and ΦT7 (*Podoviridae*). Phages were chosen due to their absence in soil and as representatives of the three families belonging to the dsDNA most-abundant bacteriophage family: *Caudovirales*. Enterobacteria phage T7 and *Listeria* phage A511 were obtained from our in-house stock. *Staphylococcus* phage 2638AΔLCR is a modified version of phage 2638A lacking the lysogenic control region (LCR) (Samuel Kilcher, unpublished), and was chosen to assess the rate of recovery of phages infecting this bacterial genus.

### 16S rRNA gene qPCR analysis

The presence of contaminating bacterial DNA was assessed throughout each extraction route (Additional file 1: Figure S1) using Taqman 16S rRNA qPCR. For this, a 16S rRNA gene fragment (934 bp) was cloned into a pGEM-T-Easy-Vector (3,015 bp), and the correct insertion was verified using restriction enzyme digestion. For standard preparation, the plasmid containing the insert was linearized and purified with Gene-Elute PCR Clean-Up Kit (Sigma). DNA concentration was measured with Qubit (ThermoFisher) and copy numbers were calculated. Taqman qPCR was carried out using the SensiFAST Probe no-ROX kit (Bioline), whereas primer and probes were placed in conserved regions of the 16S rRNA gene (amplicon size: 105 bp, Additional file 2: Table S1). All qPCR assays were performed on Rotorgene 600 (BioLabo, Corbett Research) with following conditions: 5 minutes at 95 °C for polymerase activation, followed by 40 cycles at 95 °C for 10 sec and 60 °C for 20 sec.

### Plaque assay

Spiked bacteriophage recovery was quantified at each of the optimization steps (Additional file 1: Figure S1), using plaque assays. ΦA511, Φ2638AΔLCR, and ΦT7 were titrated on host strains *Listeria ivanovii* WSLC 3009, *Staphylococcus aureus* 2638A, and *Escherichia coli* DSM496, respectively. All plaque assays were carried out using LC agar as top agar (10 g/L casein pepton (LLG), 5 g/L yeast extract (LLG), 128 mM NaCl, 55.5 mM glucose, 2mM MgSO_4,_ 10 mM CaCl_2_, 0.4 % agar). As bottom agar, brain heart infusion agar (18.5 g/L, 2 % agar, Biolife) was used for *L. ivanovii* 3009 and *S. aureus* 2638A, and Luria-Bertani agar (per litre, 10 g casein peptone, 5 g yeast extract, 7 g NaCl, 2 % agar, pH 7.2) for *E. coli* DSM496. Phages (10 µL, serial diluted) were mixed with hosts in molten soft agar (47 °C), and plates were incubated for 16 hours prior quantification at 37 °C for *E. coli* DSM496 and *S. aureus* 2638A, and at 30 °C for *L. ivanovii* 3009. The absence of spiked phages (100 µL, undiluted) in each soil sample was confirmed by plaque assay from eluted soil on all host strains.

### Elution of bacteriophages from soil

The elution optimization strategy is summarized in Figure 1. Soil samples (400 g) were spiked for each phage with 1 × 10^6^ pfu g^−1^ soil. The spiked sample was suspended 1:1 (w/v) in the respective elution buffer and manually shaked for 10 minutes by repetitive inversion. Elution buffers previously proposed in literature, such as SM buffer [9, 10] (200 mM sodium chloride, 10 mM MgSO_4_, 50 mM tris and 0.01 % gelatin, pH 7.4), 10 % beef extract buffer (10 % beef extract in ddH_2_O) [3, 18], PBS supplemented with beef extract [32] (PBS, 2.5 % beef extract, pH 8.5) and AKC [1] (1 % KC, 10 % PBS, 150 mM MgSO_4_, 5 mM EDTA) were tested for efficiency. Those elution buffers were additionally altered in constitution and assessed for efficacy with the following modifications: SM buffer was supplemented with 0.01 % tween or reduced in magnesium (5 mM MgSO_4_), and elution buffers PPBS (2 % BSA, 10 % PBS, 1 % KC, 150 mM MgSO_4_) and BPBS (2 % beef extract, 10 % PBS, 1 % KC, 150 mM MgSO_4_) were created. After the gentle elution, soil samples were left in suspension overnight at 4 °C. The next day, suspended soil samples were either directly subjected to bacterial DNA elimination (Figure 1, Route 1-4), or elsewise, remaining soil pellets were resuspended twice more (Figure 1, Route 5-6). For this, the eluted overnight soil sample was centrifuged 10,000 x g [4, 10], 10 minutes at 4°C and the first supernatant kept aside. Consecutively, the same soil pellet was again suspended in equal parts of the buffer, put on a shaker for 30 min at 300 rpm, 4°C, and centrifuged as described above. This was repeated a third time to maximise bacteriophage recovery [4, 12]. PFU for each spiked phage was assessed in all supernatants and the three finally united.

### Removal of bacterial contamination

In order to reduce contaminating bacteria and sediments, the single (Figure 1, Route 1-4) or united (Figure 1, Route 5-6) supernatants were centrifuged three rounds at 5,000 x g [20], for 10 minutes at 4 °C. At each individual round, the supernatant was recovered into a new, sterile centrifugation tube and the pellet discarded. To remove larger floating particles that were not parted by centrifugation, soil supernatants were pre-filtrated using a 16 µm cellulose filter and sterile glassware. The filtrate was eventually passed through either a 0.45 µm or 0.22 µm PES filter. Bacterial contamination, as well as recovered bacteriophages, were determined using 16S rRNA gene qPCR and plaque assays, respectively. Besides centrifugation and filtration, the efficiency of chloroform treatment to remove bacterial contamination was assessed. To evaluate potential benefits of chloroform and to concurrently allow downstream concentration of viral particles, a final chloroform concentration of 0.8 % was applied (Figure 1, Route 1-2).

### Tangential flow filtration

For concentrating soil viral particles, a TFF approach was tested. Briefly, viral suspensions were concentrated using a 100 kDa cut-off PES membrane (Millipore) and the retentate containing the bacteriophages (>100 kDa) was continuously cycled to maximally reduce its volume. The presence of spiked bacteriophages was quantified in the retentate and their absence confirmed in the permeate. TFF concentrated soil viral suspensions were either purified with CsCl ultracentrifugation, or directly subjected to DNA extraction.

### Polyethylene glycol precipitation

Concentration of soil viral particles using PEG precipitation was performed as following. Soil suspensions were mixed thoroughly 2:1 with a 3 X precipitant solution (30 % PEG 6,000 and 3 M NaCl in autoclaved ddH_2_O), and put in ice-water at 4 °C overnight. Next, the suspensions were centrifuged at 16,000 x g for 1 hour at 4 °C and the pellet resuspended in 7 mL of SM buffer. To free the concentrated viral particles of PEG, samples were dialyzed in 4 litres of SM buffer overnight at RT using a 50 kDa membrane (Biotech CE tubing).

### Caesium chloride gradient purification

Concentrated viral particles were purified using ultracentrifugation in a four-layered CsCl gradient. Methods were adapted from literature [9, 20]. Briefly, 10 mL of a concentrated sample was adjusted to a density of 1.15 g mL^−1^ and loaded on top of a 6 mL step gradient containing 2 mL of 1.35, 1.5 and 1.7 g mL^−1^ CsCl, respectively. Gradients were centrifuged at 82,000 x g for 2 hours at 10 °C. Bacteriophages in the density fractions between 1.35 and 1.5 were harvested (position visible through a light blue phage band). The collected samples with a finale volume of 2-3 mL per gradient were dialyzed at 4 °C as described above.

### Ultrafiltration concentration

The TFF retentate and dialyzed CsCl fractions were further concentrated using Amicon ultra centrifugal filters with 100 kDa cut-off (Millipore). Prior to centrifugation, filters were coated with PBS + 2 % BSA in order to prevent viral absorption [27]. The concentrated sample was recovered into a sterile Eppendorf and the centrifugal filter washed twice with 100 µL of ddH_2_O. Bacteriophage recovery in the concentrate and bacteriophage absence in the filtrate was confirmed using plaque assay. The total volume was eventually brought up to 450 µL for DNA extraction.

### Viral DNA extraction

The optimization strategy for viral DNA extraction is summarized in Figure 1 (Route A-E). Phenol chloroform viral DNA extraction was carried out as described elsewhere [9] with some optimizations. Ultrafiltrated concentrated samples were supplemented with 50 µL of 10 x DNase I Buffer (ThermoFisher) and treated with 10 U of DNase I (ThermoFisher) for 2 hours at 37 °C. The enzyme was inhibited using 50 mM EDTA at 65 °C for 10 minutes and the volume brought up to 600 µL. From here, viral DNA was either extracted using modified phenol/chloroform extraction routes (Figure 1, Route A-D) or QIAamp Viral RNA Mini Kit [22] (Qiagen) according manufacturer’s instructions (Figure 1, Route E).

For phenol/chloroform viral DNA extraction, the volume was split into two Eppendorf. Eppendorf 1 (Route A-B) was treated, following recommendations in Thurber et al [9] as following: 0.1 volumes of 2 M Tris HCL / 0.2 M EDTA, 1 volume of formamide and 1 µL glycogen at 20 mg/mL were added to each sample. Straight after an incubation at RT for 30 minutes, the DNA was spun down by adding 2 volumes of 99.9 % ethanol and centrifuged at 14,000 x g for 20 minutes at 4 °C. The pellet was washed twice using 70 % icecold ethanol and resuspended overnight in 300 µL of TE buffer (10 mM Tris, 0.5 M EDTA, pH 8) at 4 °C. The remaining suspension was again split into two equal volumes. Eppendorf 2, which was not treated with formamide, was equally split (Route C-D). All four Eppendorf tubes were topped up to 567 µL using sterile ddH_2_O and the DNA was extracted as following: 30 µL of 10 % SDS and 3 µL of 20 mg/mL Proteinase K were added, mixed, and incubated for 1 h at 55 °C. Subsequently, in one Eppendorf originating from the formamide treatment (Route A), and one without formamide treatment (Route C), 80 µL of CTAB/NaCl solution was added and incubated for 10 minutes at 65 °C. Finally, all samples were identically treated in accordance to established protocols [9]: equal volumes of chloroform were added, and the samples were centrifuged for 5 minutes at 8,000 x g at RT. The supernatant of each route was transferred to a separate tube and equal volumes of first phenol/chloroform/isoamyl alcohol (25:24:1) and subsequently chloroform were added, to be centrifuged at the same conditions as above. After the second chloroform treatment the supernatant was recovered, and 0.7 volumes of isopropanol were supplemented to precipitate the DNA overnight at 4 °C. The next day, all samples were centrifuged for 15 minutes at 13,000 x g, 4 °C, and the pellet was washed with 500 µL of 70 % ice-cold ethanol. The ethanol was then removed, the pellet air-dried and resuspended in 50 µL of ddH_2_O overnight. This DNA extraction optimization protocol was applied to larger volumes if the sample was not ultracentrifuged but directly originated from TFF concentration.

### Library preparation, illumina sequencing and annotation

The four most optimal extraction routes, e.g. 0.22 μm + TFF, 0.22 μm + PEG, 0.45 μm + TFF and 0.45 μm + PEG, were selected based on spiked phage recovery and bacterial depletion. Those extraction protocols were then used to extract viral DNA originating from 1 Kg of freshly agricultural soil (ZOFE, see above) and shotgun sequenced. For this, soil samples were suspended in PPBS, filtrated, concentrated, and purified with CsCl ultracentrifugation. Viral DNA was obtained using the optimized DNA extraction protocol including CTAB but neglecting formamide. Libraries were prepared with 25 ng unamplified viral DNA of each respective route using NebNext Ultra II DNA Library Prep for Illumina and following the manufacturer’s instructions (10 rounds of PCR amplification). Library pooling and normalization was based on the concentration of the final libraries as determined with Tapestation (Agilent 4200). Tagged libraries were sequenced with 76 million paired-end reads (150 bp/read) using NextSeq 500 sequencing. Raw reads were trimmed with Trimmomatic in default settings and unshuffled trimmed reads paired into a single file using shuffleSequences [33, 34]. Shuffled sequences from each individual metavirome were assembled with IDBA-UD [35] and contigs larger than 5 kb were extracted for further analysis. Open reading frames (ORFs) on assembled contigs (> 5 kb) were predicted using prodigal [36] and annotated using DIAMOND [37] against the NCBI NR, and using HMMscan [38] against the Pfam [39], COGs [40] and TIGRfams [41] databases.

### 16S rRNA gene contamination

In order to assess 16S rRNA gene contamination in the four sequenced metaviromes, reads from each sample were trimmed to a minimum length of 50 bp, and a random subset of 20 million trimmed reads were kept for further analysis. Potential 16S rRNA DNA reads were retrieved by USEARCH6 [42] against the RDP database [43], previously clustered at 90 % identity [43]. The percentage of 16S rRNA gene reads in each method was then calculated based on hits that were confirmed by ssu-align [44]. Affirmed 16S rRNA gene reads were taxonomically classified using the RDP database and classifications with a sequence match (S_ab score) higher than 0.8 were kept. In order to compare the efficiency of bacterial DNA removal with the here optimized protocols, 16S rRNA gene reads originating from a metavirome extracted from the same soil but using a former standardised protocol published in literature (LIT) [9, 20], was also processed as described above.

### Classification of contigs and cluster analysis for complete viral genomes

In a first step, annotated contigs (> 5 kb) from the four sequenced metaviromes were manually inspected and classified. Contigs were assigned as viral if phage structural genes such as terminase, portal, capsid or tail proteins were present, or if the majority of taxonomical hits belonged to virus. Elsewise, contigs with ribosomal proteins, cell division proteins or other bacteria hallmark proteins were classified as bacteria. Contigs with no evident gene indicators or contigs with proteins of none or hypothetical functions were left as unclassified [21]. Manually assigned viral contigs were then pooled together and viral redundant sequences removed. For this, all viral contigs were globally aligned [45] and clustered at > 99 % identity, whereas only the largest representative contig of each cluster was kept. Phage genomes were then assessed for completeness by searching for overlapping nucleotide sequences (> 10 bp) at the 3′ and 5′ region.

### Viral recruitment comparison in phage extraction routes

Viral diversity in each extraction route was compared by mapping a subset of 20 million reads from all four metaviromes to the total extracted soil viral community. This viral community originated from the manually curated viral contigs that survived the first and a second cluster round. For the second cluster analysis, remaining viral contigs were anew clustered, if more than 30 % of a smaller contig was present at > 99 % identity in a larger contig (local alignment) [45]. Reads of each metavirome were then mapped against all viral contigs that were cleared for redundant sequences, and phage abundance and diversity from each optimized method was analysed. To provide a normalized measure, the number of hits to each phage contig was divided by the length of the contig (in kb) and by the size of the metavirome (size of the database in Gb). This measure is abbreviated as RPKG (reads per Kb per Gb) and helps to compare recruitments by differently sized contigs versus several metagenomes. A phage was considered present in a given metavirome, if the contig was covered by at least by 1 RPKG at 98 % identity.

### Metavirome reads associated with bacterial, viral or unclassified contigs

A subset of 20 million reads of each extraction route was mapped to the manually classified viral, bacterial or unknown contigs. For this, all viral sequences obtained from the extracted metaviromes were concatenated to one super viral DNA contig. All bacteria or unknown sequences were likewise combined. This concatenation prevents multiple mapping of a single read to a given viral sequence if represented several times in the assembled metaviromes. The total percentage of reads recruited to either the phage, bacterial or unclassified concatenated super contig was then assessed at 98 % identity.

## Abbreviations

Phages: bacteriophages
PEG: polyethylene glycol
TFF: tangential flow filtration
CsCl: caesium chloride
PBS: phosphate buffered saline
AKC: amended potassium citrate
RT: room temperature
BSA: bovine serum albumin
PPBS: protein supplemented PBS
BPBS: beef extract supplemented PBS.

## Declaration

### Acknowledgments

We thank Jochen Mayer and Lucie Gunst (Agroscope, Zürich) for giving us the opportunity to take soil samples from the ZOEFE experimental field, their help during soil sampling and for providing the physiochemical parameters of the soil samples at the date of sampling. We further would like to thank Dr. Samuel Kilcher (ETH Zurich) for providing us with phage 2638AΔLCR.

### Availability of data and materials

Metagenomic datasets have been submitted to NCBI SRA and are available under BioProject accession number PRJNA544697 (ZOFE18-TFF_0.22 [SAMN11866222], ZOFE18-TFF_0.45 [SAMN11866225], ZOFE18-PEG_0.22 [SAMN11866227], ZOFE18-PEG_0.45 [SAMN11866228] and ZOFE16-VIR-MAN [SAMN11866229]).

### Author’s contribution

EGS conceived the study, helped to design experiments, and contributed to the writing of the manuscript. PCG designed, performed, and analysed all experiments and wrote the manuscript. JMH and FRV planned and guided the sequencing analysis of all sequenced metaviromes and corrected the manuscript. MJL guided all experimental work, helped to analyse results and corrected the manuscript. All authors read and approved the finale manuscript.

## Additional files

### Additional file 1

Figure S1: Optimization Strategy (PDF)

Optimization strategy of phage extractions protocols form soil samples prior to metagenomics analysis. Different phage elution, filtration, concentration and DNA extraction procedures were tested to maximise viral yield and deplete bacterial DNA contaminates. *16S rRNA qPCR to determine external contaminates, ^I^ plaque assay to assess spiked bacteriophage recovery.

### Additional file 2

Table S1: Primer and probes used for 16S rRNA qPCR (PDF) Primer and probes designed for TaqMan 16S rRNA gene qPCR.

### Additional file 3

Table S2: DNA yield and external contamination with phage DNA extraction methods (PDF) DNA yield and bacterial DNA contamination obtained from phage DNA extraction of soil.

### Additional file 4

Table S3: Assembly with 60 million reads (PDF)

Normalized viral metagenomic assembly with 60 million reads.

## References

1. Trubl G, Solonenko N, Chittick L, Solonenko SA, Rich VI, Sullivan MB: Optimization of viral resuspension methods for carbon-rich soils along a permafrost thaw gradient. PeerJ 2016, 4:e1999.

2. Pratama AA, van Elsas JD: The ‘Neglected’ Soil Virome - Potential Role and Impact. Trends Microbiol 2018.

3. Williamson KE, Wommack KE, Radosevich M: Sampling natural viral communities from soil for culture-independent analyses. Appl Environ Microbiol 2003, 69:6628–6633.

4. Williamson KE, Radosevich M, Wommack KE: Abundance and diversity of viruses in six Delaware soils. Appl Environ Microbiol 2005, 71:3119–3125.

5. Amossé J, Bettarel Y, Bouvier C, Bouvier T, Tran Duc T, Doan Thu T, Jouquet P: The flows of nitrogen, bacteria and viruses from the soil to water compartments are influenced by earthworm activity and organic fertilization (compost vs. vermicompost). Soil Biol Biochem 2013, 66:197–203.

6. Cobian Guemes AG, Youle M, Cantu VA, Felts B, Nulton J, Rohwer F: Viruses as Winners in the Game of Life. Annu Rev Virol 2016, 3:197–214.

7. Adriaenssens EM, Van Vaerenbergh J, Vandenheuvel D, Dunon V, Ceyssens PJ, De Proft M, Kropinski AM, Noben JP, Maes M, Lavigne R: T4-related bacteriophage LIMEstone isolates for the control of soft rot on potato caused by ‘Dickeya solani’. PLoS One 2012, 7:e33227.

8. Zablocki O, van Zyl L, Adriaenssens EM, Rubagotti E, Tuffin M, Cary SC, Cowan D: High-level diversity of tailed phages, eukaryote-associated viruses, and virophage-like elements in the metaviromes of antarctic soils. Appl Environ Microbiol 2014, 80:6888–6897.

9. Thurber RV, Haynes M, Breitbart M, Wegley L, Rohwer F: Laboratory procedures to generate viral metagenomes. Nat Protoc 2009, 4:470–483.

10. Casas V, Rohwer F: Phage Metagenomics. Methods Enzymol 2007, 421:259–268.

11. Narr A, Nawaz A, Wick LY, Harms H, Chatzinotas A: Soil Viral Communities Vary Temporally and along a Land Use Transect as Revealed by Virus-Like Particle Counting and a Modified Community Fingerprinting Approach (fRAPD). Front Microbiol 2017, 8.

12. Swanson MM, Fraser G, Daniell TJ, Torrance L, Gregory PJ, Taliansky M: Viruses in soils: morphological diversity and abundance in the rhizosphere. Ann Appl Biol 2009, 155:51–60.

13. Quiros P, Muniesa M: Contribution of cropland to the spread of Shiga toxin phages and the emergence of new Shiga toxin-producing strains. Sci Rep 2017, 7:7796.

14. Williamson KE, Radosevich M, Smith DW, Wommack KE: Incidence of lysogeny within temperate and extreme soil environments. Environ Microbiol 2007, 9:2563–2574.

15. Adriaenssens EM, Van Zyl L, De Maayer P, Rubagotti E, Rybicki E, Tuffin M, Cowan DA: Metagenomic analysis of the viral community in Namib Desert hypoliths. Environ Microbiol 2015, 17:480–495.

16. Ashelford KE, Day MJ, Fry JC: Elevated Abundance of Bacteriophage Infecting Bacteria in Soil. Appl Environ Microbiol 2003, 69:285–289.

17. Reavy B, Swanson MM, Cock PJ, Dawson L, Freitag TE, Singh BK, Torrance L, Mushegian AR, Taliansky M: Distinct circular single-stranded DNA viruses exist in different soil types. Appl Environ Microbiol 2015, 81:3934–3945.

18. Lasobras J, Dellunde J, Jofre J, Lucena F: Occurrence and levels of phages proposed as surrogate indicators of enteric viruses in different types of sludges. J Appl Microbiol 1999, 86:723–729.

19. Kimura M, Jia ZJ, Nakayama N, Asakawa S: Ecology of viruses in soils: Past, present and future perspectives. Soil Sci Plant Nutr 2008, 54:1–32.

20. Castro-Mejia JL, Muhammed MK, Kot W, Neve H, Franz CM, Hansen LH, Vogensen FK, Nielsen DS: Optimizing protocols for extraction of bacteriophages prior to metagenomic analyses of phage communities in the human gut. Microbiome 2015, 3:64.

21. Enault F, Briet A, Bouteille L, Roux S, Sullivan MB, Petit MA: Phages rarely encode antibiotic resistance genes: a cautionary tale for virome analyses. ISME J 2017, 11:237–247.

22. Conceicao-Neto N, Zeller M, Lefrere H, De Bruyn P, Beller L, Deboutte W, Yinda CK, Lavigne R, Maes P, Van Ranst M, et al: Modular approach to customise sample preparation procedures for viral metagenomics: a reproducible protocol for virome analysis. Sci Rep 2015, 5:16532.

23. Hoyles L, McCartney AL, Neve H, Gibson GR, Sanderson JD, Heller KJ, van Sinderen D: Characterization of virus-like particles associated with the human faecal and caecal microbiota. Res Microbiol 2014, 165:803–812.

24. Fierer N, Breitbart M, Nulton J, Salamon P, Lozupone C, Jones R, Robeson M, Edwards RA, Felts B, Rayhawk S, et al: Metagenomic and small-subunit rRNA analyses reveal the genetic diversity of bacteria, archaea, fungi, and viruses in soil. Appl Environ Microbiol 2007, 73:7059–7066.

25. Zablocki O, Adriaenssens EM, Cowan D: Diversity and Ecology of Viruses in Hyperarid Desert Soils. Appl Environ Microbiol 2016, 82:770–777.

26. Subirats J, Sanchez-Melsio A, Borrego CM, Balcazar JL, Simonet P: Metagenomic analysis reveals that bacteriophages are reservoirs of antibiotic resistance genes. Int J Antimicrob Agents 2016, 48:163–167.

27. Deng L, Gregory A, Yilmaz S, Poulos BT, Hugenholtz P, Sullivan MB: Contrasting Life Strategies of Viruses that Infect Photo- and Heterotrophic Bacteria, as Revealed by Viral Tagging. Mbio 2012, 3.

28. Wang Q, Garrity GM, Tiedje JM, Cole JR: Naive Bayesian classifier for rapid assignment of rRNA sequences into the new bacterial taxonomy. Appl Environ Microbiol 2007, 73:5261–5267.

29. He X, McLean JS, Edlund A, Yooseph S, Hall AP, Liu SY, Dorrestein PC, Esquenazi E, Hunter RC, Cheng G, et al: Cultivation of a human-associated TM7 phylotype reveals a reduced genome and epibiotic parasitic lifestyle. Proc Natl Acad Sci U S A 2015, 112:244–249.

30. Paez-Espino D, Chen IA, Palaniappan K, Ratner A, Chu K, Szeto E, Pillay M, Huang J, Markowitz VM, Nielsen T, et al: IMG/VR: a database of cultured and uncultured DNA Viruses and retroviruses. Nucleic Acids Res 2017, 45:D457–D465.

31. Paez-Espino D, Eloe-Fadrosh EA, Pavlopoulos GA, Thomas AD, Huntemann M, Mikhailova N, Rubin E, Ivanova NN, Kyrpides NC: Uncovering Earth’s virome. Nature 2016, 536:425–430.

32. Quirós P, Maryury BJ, Maite M: Spread of bacterial genomes in packaged particles. Future Microbiol 2016, 11:171–173.

33. Bolger AM, Lohse M, Usadel B: Trimmomatic: a flexible trimmer for Illumina sequence data. Bioinformatics 2014, 30:2114–2120.

34. Zerbino DR, Birney E: Velvet: Algorithms for de novo short read assembly using de Bruijn graphs. Genome Res 2008, 18:821–829.

35. Peng Y, Leung HC, Yiu SM, Chin FY: IDBA-UD: a de novo assembler for single-cell and metagenomic sequencing data with highly uneven depth. Bioinformatics 2012, 28:1420–1428.

36. Hyatt D, Chen GL, LoCascio PF, Land ML, Larimer FW, Hauser LJ: Prodigal: prokaryotic gene recognition and translation initiation site identification. BMC Bioinformatics 2010, 11.

37. Buchfink B, Xie C, Huson DH: Fast and sensitive protein alignment using DIAMOND. Nat Methods 2015, 12:59–60.

38. Eddy SR: Accelerated Profile HMM Searches. PLoS Comput Biol 2011, 7:e1002195.

39. Bateman A, Coin L, Durbin R, Finn RD, Hollich V, Griffiths-Jones S, Khanna A, Marshall M, Moxon S, Sonnhammer EL, et al: The Pfam protein families database. Nucleic Acids Res 2004, 32:D138–141.

40. Tatusov RL, Fedorova ND, Jackson JD, Jacobs AR, Kiryutin B, Koonin EV, Krylov DM, Mazumder R, Mekhedov SL, Nikolskaya AN, et al: The COG database: an updated version includes eukaryotes. BMC Bioinformatics 2003, 4:41.

41. Haft DH, Loftus BJ, Richardson DL, Yang F, Eisen JA, Paulsen IT, White O: TIGRFAMs: a protein family resource for the functional identification of proteins. Nucleic Acids Res 2001, 29:41–43.

42. Edgar RC: Search and clustering orders of magnitude faster than BLAST. Bioinformatics 2010, 26:2460–2461.

43. Cole JR, Wang Q, Fish JA, Chai B, McGarrell DM, Sun Y, Brown CT, Porras-Alfaro A, Kuske CR, Tiedje JM: Ribosomal Database Project: data and tools for high throughput rRNA analysis. Nucleic Acids Res 2014, 42:D633–642.

44. Nawrocki EP, Eddy SR: Infernal 1.1: 100-fold faster RNA homology searches. Bioinformatics 2013, 29:2933–2935.

45. Li W, Godzik A: Cd-hit: a fast program for clustering and comparing large sets of protein or nucleotide sequences. Bioinformatics 2006, 22:1658–1659.

