## Supplemental files for "Uncovering a hidden diversity: optimized protocols for the extraction of bacteriophages from soil"

### Additional file 1: Figure S1

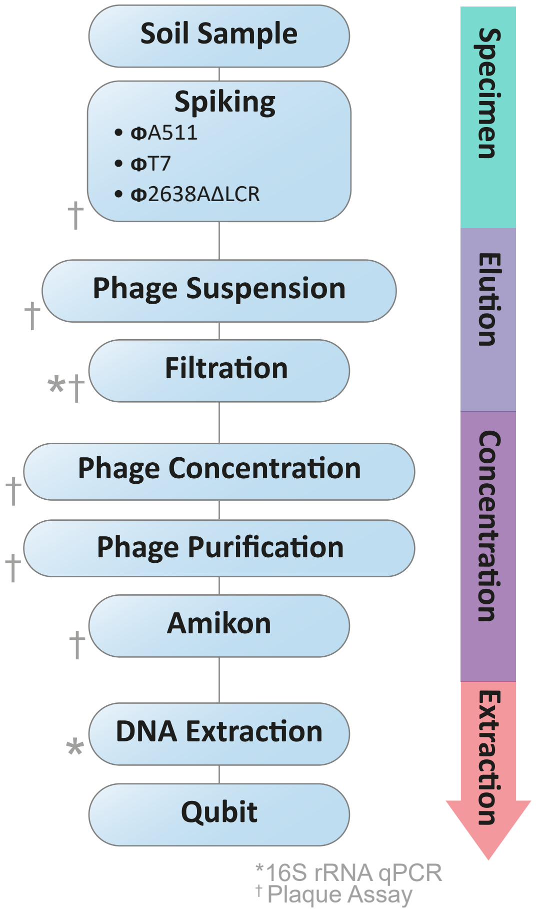


Figure S1. Optimization strategy of phage extractions protocols form soil samples prior to metagenomics analysis. Different phage elution, filtration, concentration and DNA extraction procedures were tested to maximise viral yield and deplete bacterial DNA contaminates. *16S rRNA qPCR to determine external contaminates, ł plaque assay to assess spiked bacteriophage recovery.

### Additional file 2: Table S1

Table S1. Primer and probes designed for TaqMan 16S rRNA gene qPCR.

| **Oligos** | **Sequence (5'-3')** |
|---|---|
| Primer_fw | GCGGTGAAATGCGTAGAGAT |
| Primer_rv | TCTAATCCTGTTTGCTCCCCA |
| Probe | FAM-GCGAAGGCGGCCCCCTGGAC-BHQ1 |

### Additional file 3: Table S2

Table S2. DNA yield and bacterial DNA contamination obtained from phage DNA extraction of soil.

| CsCl Purified Sample | 16S rRNA (copies/μL) | Inhibition | DNA in 400 g of soil (ng/μL) |
|---|---|---|---|
| No Formamide, no CTAB | 2.22E+03 | - | 12.4 |
| No Formamide, CTAB | 3.73E+02 | - | 24.4 |
| Formamide, no CTAB | 1.81E+03 | - | 10.2 |
| Formamide, CTAB | 1.56E+03 | - | 6.2 |

### Additional file 4: Table S3

Table S3. Normalized viral metagenomic assembly with 60 million reads.

| Sample | Trimmed Reads (Mill) | Contigs (> 5 kb) | Nucleotides Assembled (Mb) |
|---|---|---|---|
| 0.22 μm TFF | 60 | 11,215 | 125.1 |
| 0.22 μm PEG | 60 | 7,577 | 84.5 |
| 0.45 μm TFF | 60 | 9,517 | 110.5 |
| 0.45 μm PEG | 60 | 9,288 | 117.5 |
